# The expression of hemagglutinin by a recombinant Newcastle disease virus causes structural changes and alters innate immune sensing

**DOI:** 10.1101/2020.02.19.957084

**Authors:** Ingrao Fiona, Duchatel Victoria, Fernandez Rodil Isabel, Steensels Mieke, Verleysen Eveline, Mast Jan, Lambrecht Bénédicte

**Author notes:** Corresponding author: Fiona Ingrao.

## Abstract

Recombinant Newcastle disease viruses (rNDV) have been used as bivalent vectors for vaccination against multiple economically important avian pathogens. NDV-vectored vaccines expressing the immunogenic H5 hemagglutinin (rNDV-H5) are considered attractive candidates to protect poultry from both highly pathogenic avian influenza (HPAI) and Newcastle disease (ND). However, the impact of the insertion of a recombinant protein, such as H5, on the biological characteristics of the parental NDV strain has been little investigated to date. The present study compared a rNDV-H5 vaccine and its parental NDV LaSota strain in terms of their structural and functional characteristics, as well as their recognition by the innate immune sensors. Structural analysis of the rNDV-H5 demonstrated a decreased number of fusion (F) and a higher number of hemagglutinin-neuraminidase (HN) glycoproteins compared to NDV LaSota. These structural differences were accompanied by increased hemagglutinating and neuraminidase activities of rNDV-H5. During *in vitro* rNDV-H5 infection, increased mRNA expression of TLR3, TLR7, MDA5, and LGP2 was observed, suggesting that the recombinant virus is recognized differently by sensors of innate immunity when compared with the parental NDV LaSota. Given the growing interest in using NDV as a vector against human and animal diseases, these data highlight the importance to thoroughly understand the recombinant vaccines structural organization, functional characteristics, and the immune responses elicited.

## INTRODUCTION

Newcastle disease (ND) and highly pathogenic avian influenza (HPAI) are two highly contagious and economically devastating notifiable poultry diseases. The continuous threat they represent to the poultry sector worldwide emphasizes the need for high level of biosecurity measures, strong surveillance strategies, and efficient vaccination programs [1–3]. Both attenuated and inactivated vaccines have been extensively and successfully used to protect poultry from infectious diseases, but they can also have drawbacks that have been previously reviewed [4,5]. The development of new vaccination strategies is an answer to the call for more protective vaccines that would overcome the challenges faced by classical immunization. The use of recombinant viral vector-based vaccines expressing one or several foreign genes represents a promising vaccination strategy. Live recombinant vaccines have the advantage of eliciting cellular and mucosal responses as well as humoral immunity. Herpesvirus of turkey, NDV LaSota, and fowlpox viruses are widely used veterinary vaccine virus backbones [6] and NDV is also considered as promising viral vector vaccine candidate against human diseases [7]. The protection afforded by these vaccines is validated by testing their potency following an infection by a virulent pathogen related to the viral vector or the foreign gene expressed, and by identifying the host immune responses elicited. However, a shortcoming of this approach is the lack of systematic investigation of the impact of foreign gene insertion into the genome of the vector on its structure and biological functions. Recombinant NDV vectored vaccines expressing the protective antigen H5 (rNDV-H5) have demonstrated their efficacy to protect specific pathogens free (SPF) chickens against both homologous and heterologous HPAI H5 and velogenic ND challenges [8–12]. NDV infection is initiated by the attachment of the virus through the binding of hemagglutinin-neuraminidase (HN) glycoprotein to the sialic-acid-receptor at the surface of the host cell [13]. As a paramyxovirus, NDV is known to enter its target cell through direct fusion with the cell membrane, and it has been suggested to use a caveolae-dependent endocytic pathway as an alternative route for viral entry [14,15]. A previous study examining a recombinant NDV expressing the glycoprotein GP of the Ebola virus showed that it used GP-dependent macropinocytosis as a major cell entry pathway, indicating that the foreign GP can function as an entry protein [16]. The AIV entry process begins with the binding of the hemagglutinin (HA) to sialic acids at the cell surface and the internalization of the viral particle by endocytosis. The low pH within the endosome triggers conformational changes in the HA, exposing the fusion peptide and inducing the fusion between the virus and the endosomal membrane [17]. A previously published study suggested the ability of rNDV-H5 to use an H5-dependent entry pathway under certain conditions, such as the presence of ND maternal antibodies [18], could therefore affect vaccine-induced immune responses. Because the latter may differ from the immune responses induced by the parental NDV LaSota, their characterization would improve the understanding of protection outcomes previously observed with rNDV-H5 immunization. In this study, the analyses focused on innate responses known to be involved in the regulation and orientation of subsequent adaptive responses [19]. Pathogen recognition by innate immune system is mediated through the sensing by pattern recognition receptors (PRRs). The activation of these receptors generates signals that trigger intracellular cascades resulting in the production of key soluble mediators that influence the polarization of adaptive immune responses [20]. Among PRRs, toll-like receptors (TLRs) −3 (TLR3) and −7 (TLR7) are important virus sensors capable of recognizing nucleic acids in intracellular compartments like endosomes. Melanoma differentiation-associated gene 5 (MDA5) and laboratory of genetics and physiology 2 (LGP2) are RNA-sensing PRRs expressed in the cytoplasm that play a key role in the activation of viral sensing pathway [21,22]. The recognition of nucleic acids derived from pathogens during an infection ultimately leads to the production of type-I interferons (IFNs) that mediates the antiviral response [23] and cytokines that influence the polarization of adaptive immune responses. To determine if the recombinant NDV-H5 retain structural and functional characteristics of the parental NDV LaSota strain, the present study compared the structural organization and enzymatic activity of surface glycoproteins, and the recognition of both viruses by innate immune sensors.

## MATERIALS AND METHODS

### Chickens

SPF White Leghorn chickens were hatched from embryonated eggs purchased from Lohmann Valo (Cuxhaven, Germany). After hatching, the chickens were housed in biosecurity level 3 isolators. Feed and water were provided *ad libitum* throughout the experimental period. The animal experiment was conducted under the authorization and supervision of the Biosafety and Bioethics Committees at Sciensano (Brussels, Belgium; bioethics authorization no. 20170413-01) according to national and European regulations.

### Vaccines and viruses

The rNDV-H5 vaccine expressing a modified H5 ectodomain of human HPAI H5N1 clade 1 A/Vietnam/1203/04, and the NDV LaSota were provided by Lohmann Animal Health GmbH (Germany) [10]. The H5 insert of the rNDV-H5 had been modified into a low-pathogenic version to ensure vaccine safety and the H5 transmembrane domain and cytoplasmic tail were replaced by those of the NDV F glycoprotein to allow surface expression [10,24]. The H5 gene was inserted between the phosphoprotein and matrix genes of the NDV genome, as it has been identified as the optimal insertion site for foreign gene expression [25]. The clade 1 HPAI H5N1 A/Crested-eagle/Belgium/01/2004 strain was isolated in 2004 in Belgium [26]. The strains used in this study were amplified by inoculation into the allantoic cavity of 9-11 day-old embryonated specific pathogen free (SPF) eggs. Five days after inoculation or at the death of the embryo, allantoic fluids were harvested and the isolates were titrated on primary chicken embryo fibroblasts (CEFs) to determine the tissue culture infectious dose (TCID50/ml) [27,28]. For immunogold electron microscopy analyses, viral strains were purified by differential centrifugation on a sucrose gradient, as previously described [29].

### Cells and Monoclonal antibodies

CEFs were cultured in complete medium composed of a mixture of Leibovitz’s L15 and McCoy’s 5A (1:1) media (Gibco, ThermoFisher Scientific, USA) supplemented with 2 % heat-inactivated fetal calf serum, 2 mM L-Glutamin, and 50 μg/ml gentamycin (Gibco) at 37°C under 5 % CO_2_. NDV F and HN glycoproteins and AIV H5 were detected using *in-house* monoclonal antibodies (mAbs) previously described: mouse anti-NDV F 1C3 (IgG1), mouse anti-NDV HN 4D6 (IgG2a), and mouse anti-AIV H5 5A1 (IgG1) [30,31].

### Immunogold electron microscopy

Glycoprotein expression on rNDV-H5 and NDV LaSota surface was evaluated by the previously described immunogold labeling method [18] with minor modifications. Briefly, pioloform carbon-coated copper grids (Agar Scientific, Stansted Essex, England) were pretreated with Alcian blue 8G (Gurr Microscopy Materials, Poole, England) solution at 1 % v/v in water for 10 min at room temperature (RT). The rNDV-H5 and NDV LaSota were diluted in PBS to a final concentration of 75 μg/ml and adsorbed onto pretreated grids for 10 min at RT. Anti-F, anti-HN, and anti-H5 mAbs at 1:50 dilution in PBS supplemented with 2 % of goat serum were then adsorbed to the grid. The number of gold particles at the surface of the virions was assessed in 50 representative virions. Images of immunogold-labeled virions were acquired on a Tecnai G2 Spirit electron microscope (FEI, Eindhoven, Netherlands) using bright-field transmission electron microscopy mode. To take in account the pleomorphism of NDV viruses [32], the surface of each virion was measured and the number of gold labels was then expressed per 55000 nm^2^ as an estimate of the number of gold particles per virion (#gold/virion). 55000 nm^2^ was determined as the mean surface of the virions.

### Virus neutralization

CEFs were seeded at a concentration of 5×10^5^ cells/ml in 96-well plates and incubated at 37°C for 24h. Two-fold serial dilutions of an initial concentration of 5 μg/ml of the mAbs were incubated with NDV LaSota or rNDV-H5 for 3h at 37°C in complete medium supplemented with 50 ng/ml L-1-tosylamido-2-phenylethylchloromethyl ketone (TPCK)-treated trypsin (Sigma Aldrich). Subsequently, CEFs monolayers were cultured with mixtures of mAbs and NDV LaSota or rNDV-H5, corresponding to a multiplicity of infection (MOI) of 0.01. After 24 h, half of the culture medium was replaced by fresh complete medium supplemented with TPCK-trypsin. CEFs were monitored daily over a 7 day period for the presence of a cytopathic effect.

### Evaluation of hemagglutinating and neuraminidase activities

The hemagglutinating activity of NDV LaSota and rNDV-H5 was evaluated by the standard hemagglutination assay [33]. Both viruses were serially two-fold diluted in triplicates from a starting titer of 10^7^ TCID50/ml and the HA titers were determined based on the lowest virus dilution at which full hemagglutination was observed.

The neuraminidase (NA) activity of NDV LaSota and rNDV-H5 was determined using the NA-fluor Influenza Neuraminidase Assay kit (Applied Biosystems, CA, USA) according to the manufacturer’s recommendations. Triplicates of two-fold serial dilutions of NDV LaSota and rNDV-H5 starting at a titer of 10^7^ TCID50/ml were analyzed and the NA activity was expressed as Relative Fluorescent Unit (RFU).

### Immunofluorescence

Immunofluorescence was performed as previously described [34]. Briefly, CEFs cultured in 6-well plates and the monolayer was infected with either NDV LaSota or rNDV-H5 at an MOI of 1 and incubated at 37°C for 1h. The medium was then replaced by fresh complete medium without antibiotics and the CEFs were incubated at 37°C for 0, 2, 6, 10, and 24 hours. NDV F protein was labeled with 1:100 1C3 mAb, followed by 1:100 FITC-conjugated sheep anti-mouse IgG as secondary antibody (F6257, Sigma Aldrich). Fluorescence was detected using a Leitz SMLUX microscope with a Leica DFC420C camera and images were analyzed with the Leica Application Suite LAS V.4 program.

### Tracheal organ cultures (TOCs) infection

Tracheas were aseptically collected from nine 12-day-old SPF chickens and washed with warm complete culture medium containing DMEM (Gibco) supplemented with 100 U/ml penicillin (Kela Pharma) and 1 mg/ml streptomycin (Sigma Aldrich). The upper part of the tracheas was dissected into 2-3 mm rings. The rings from the nine chickens were divided into three groups and cultured in pools of three per well in 1 ml of complete medium in a 12-well plate. Rings were cultured for 48 h at 39°C in 5% CO_2_ atmosphere. The culture medium was then removed and replaced with 0.5 ml of viral inoculum at the titer of 10^6^ TCID50/ml in complete culture medium. Virus adsorption was carried out for 1 hour at 39°C after which 1.5 ml of complete medium was added. The rings were collected after 0, 2, 6, 10 and 24 hours post-infection (hpi) and were stored in pools of three in 200 μl in RNAlater solution (Applied Biosystems, Lennik, Belgium) at −80°C until RNA extraction.

### CEFs infection

The CEFs were cultured and infected according to the protocol described above for immunofluorescence assay. At 0, 2, 6, 10, and 24 hpi, the medium was discarded and CEFs were detached using a solution of 0.25% Trypsin-EDTA (ThermoFisherScientific). The infected CEFs of two wells were pooled and were stored in RNA later at −80°C until analysis.

### RNA extraction and real-time reverse transcription (RT)-PCR

The RNA from infected TOCs and CEFs samples was extracted using the MagMAX-96 Total RNA Isolation kit (AM1830, Ambion, Applied Biosystems, Carlsbad, CA, USA). Synthesis of cDNA was performed using 250 ng of purified RNA using oligo(dT)_15_ primers (GoScriptTM Reverse Transcription System, A5001, Promega, Madison, WI, USA), according to the manufacturer’s instructions. The cDNA products were stored at −20°C until further use. The relative expression of TLR3, TLR7, MDA5, [35], LGP2 [36], IFNα [37], and IFNβ [38] was measured by RT-PCR, according to a previously published protocol [39]. HMBS and RPL0 [40] were selected as reference genes for normalization of RT-PCR results using the algorithm GeNorm (Biogazelle, Zwijnbeke, Belgium). Normalized gene expression was quantified as the fold change relative to the uninfected cells at time point 0 hpi according to the 2^−ΔΔCT^ method [41].

### Statistical analyses

Statistical analyses were performed using R statistical software and the results were visualized using the ggplot2 package for R [42]. Immunogold electron microscopy results were analyzed by Mann-Whitney–Wilcoxon test or Student’s paired *t*-test using permutation and, for normally distributed values, with one-way analysis of variance (ANOVA). The neuraminidase activity data were analyzed with one-way analysis of variance (ANOVA). Flow cytometry data were analyzed by ANOVA test using permutation or by Student’s paired *t*-test using permutation depending on the normality and homoscedasticity. RT-PCR data were analyzed with ANOVA and non-parametric Kruskal-Wallis test while innate immunity results on TOCs were analyzed with ANOVA only. P-value < 0.05 were considered statistically significant.

## RESULTS

### The recombinant virus expresses higher levels of HN glycoproteins and lower levels of F on its surface than the parental NDV

Immunogold electron microscopy was conducted to evaluate the expression of F and HN at the surface of rNDV-H5 and compare it to the distribution of these glycoproteins on parental NDV LaSota. Quantitative analysis of the labeling densities of F glycoprotein demonstrated a significantly lower expression at the surface of rNDV-H5 (4.3±0.3 gold/virion) when compared to NDV LaSota (8.4±0.5 gold particles/virion). In contrast, rNDV-H5 displayed a significantly higher number of HN molecules at its surface (25.8±1.4 gold particles/virion) than NDV LaSota (18.4±1.9 gold particles/virion) (**Figure 1a**). Immunogold labeling also confirmed the presence of H5 at the surface of rNDV-H5 (4.6±0.7 gold particles/virion), while a background level of H5 labeling of 0.8±0.2 gold particles/virion was detected on NDV LaSota surface.

**Figure 1.**
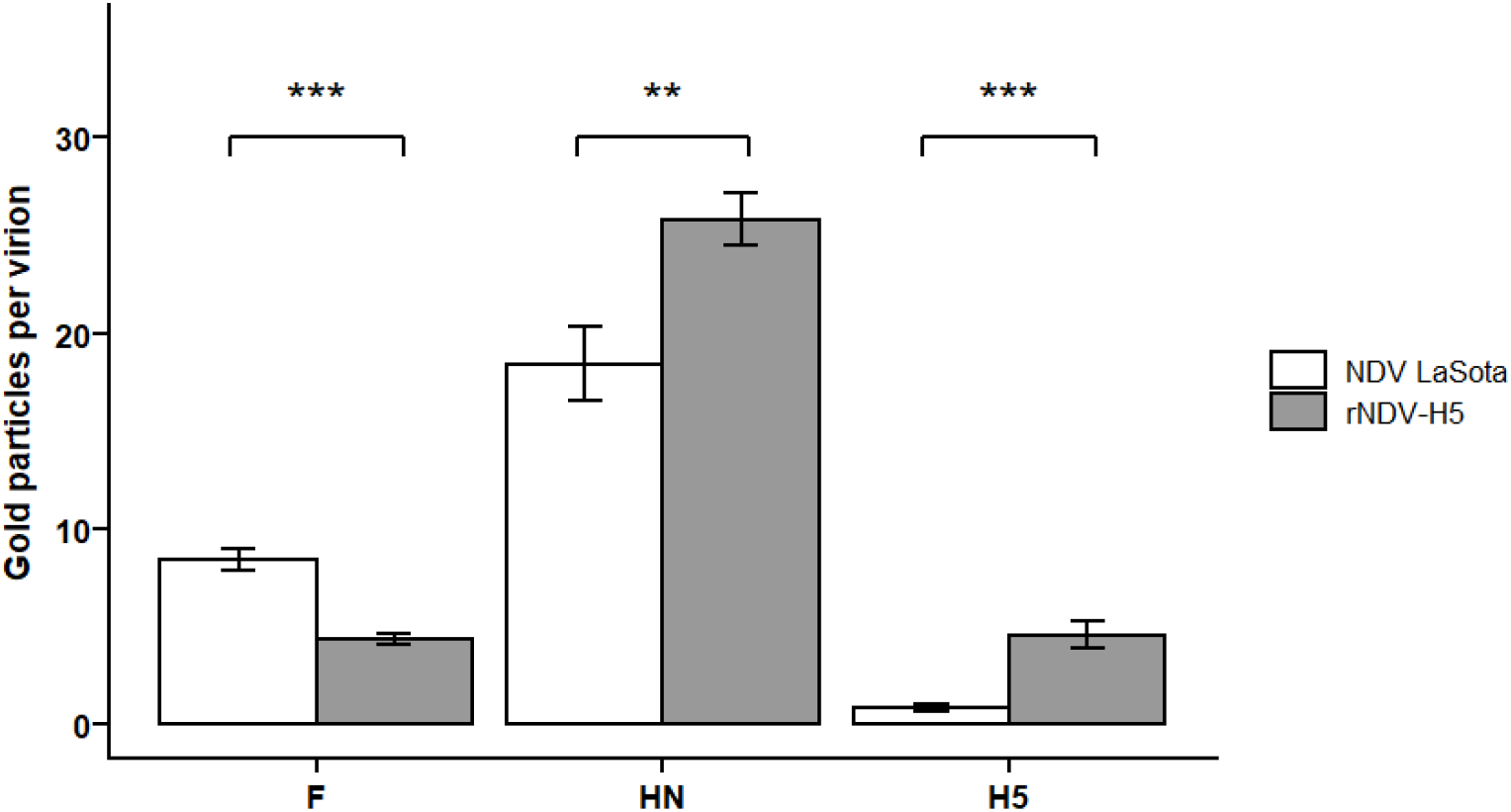
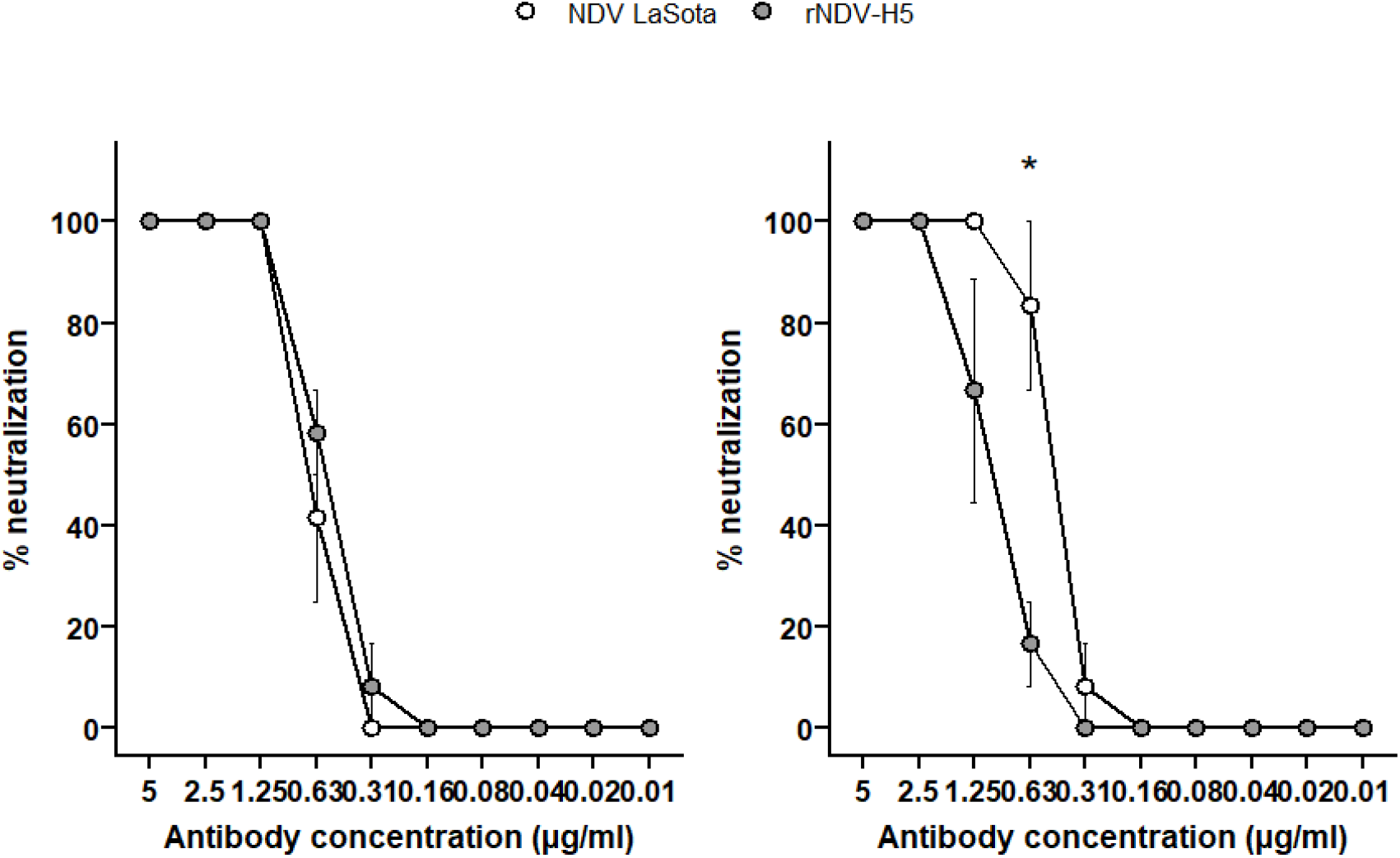
Expression of surface glycoproteins by NDV LaSota and rNDV-H5. (a) Proportion of NDV LaSota and rNDV-H5 particles expressing F, HN, and H5. Immunogold labeling of viral particles was performed with anti-HN, anti-F (1C3, IgG1), and anti-AIV H5 (5A1, IgG1) mAbs. The number of gold particles was counted and normalized per virion surface unit of 55000 nm^2^. Results are expressed as the mean ± standard error of the mean (SEM). (b) Comparison of NDV LaSota and rNDV-H5 in a neutralization test using anti-F (left panel) and anti-HN (right panel) monoclonal antibodies. Monoclonal antibodies were two-fold serially diluted and incubated with CEFs for 24h at 37°C. Percentages of neutralization are expressed for NDV LaSota (white) and rNDV-H5 (grey) according to the antibody dilutions. * p < 0.05, ** p < 0.01, ***,p < 0.001.

The capacity to block the viral entry of NDV LaSota and rNDV-H5 using anti-F and HN monoclonal antibodies was evaluated using a neutralization test. The neutralization curves obtained using the anti-F monoclonal antibody were similar for both viruses (**Figure 1b, left panel**), while NDV LaSota was neutralized at 83% by anti-HN monoclonal antibody at the concentration of 0,63 μg/ml, which was significantly higher than the 17% of neutralization of rNDV-H5 at the same mAb concentration (**Figure 1b, right panel**).

### rNDV-H5 has higher HA and NA activities than parental NDV LaSota

HA assay demonstrated that rNDV-H5 retains the ability to agglutinate chicken erythrocytes (**Figure 2a**). Hemagglutinating activity of two-fold serially diluted NDV LaSota and rNDV-H5 at the initial concentration of 10^7^ TCID50/ml was last fully detected at the titer of 6,25 × 10^5^ TCID50/ml (1:8 dilution) and 7,8 × 10^4^ TCID50/ml (1:32 dilution), respectively.

**Figure 2.**
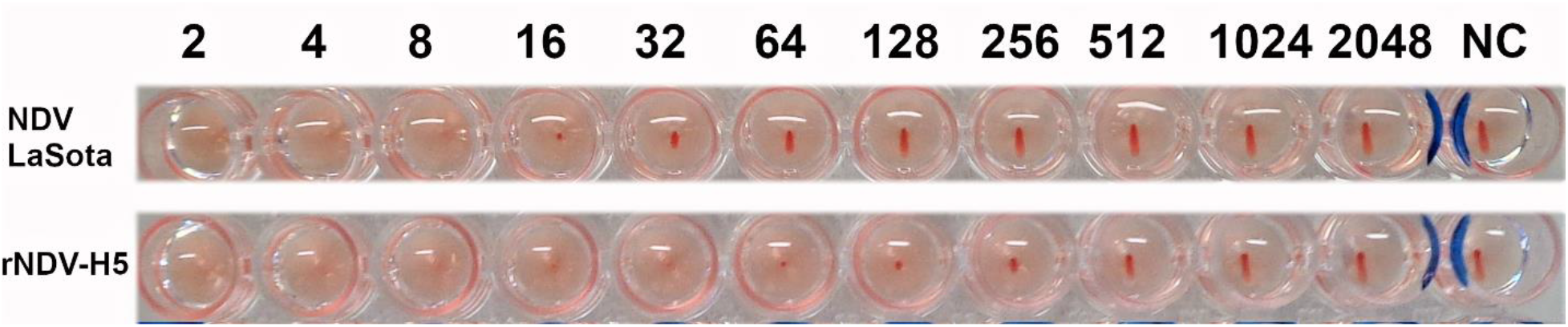
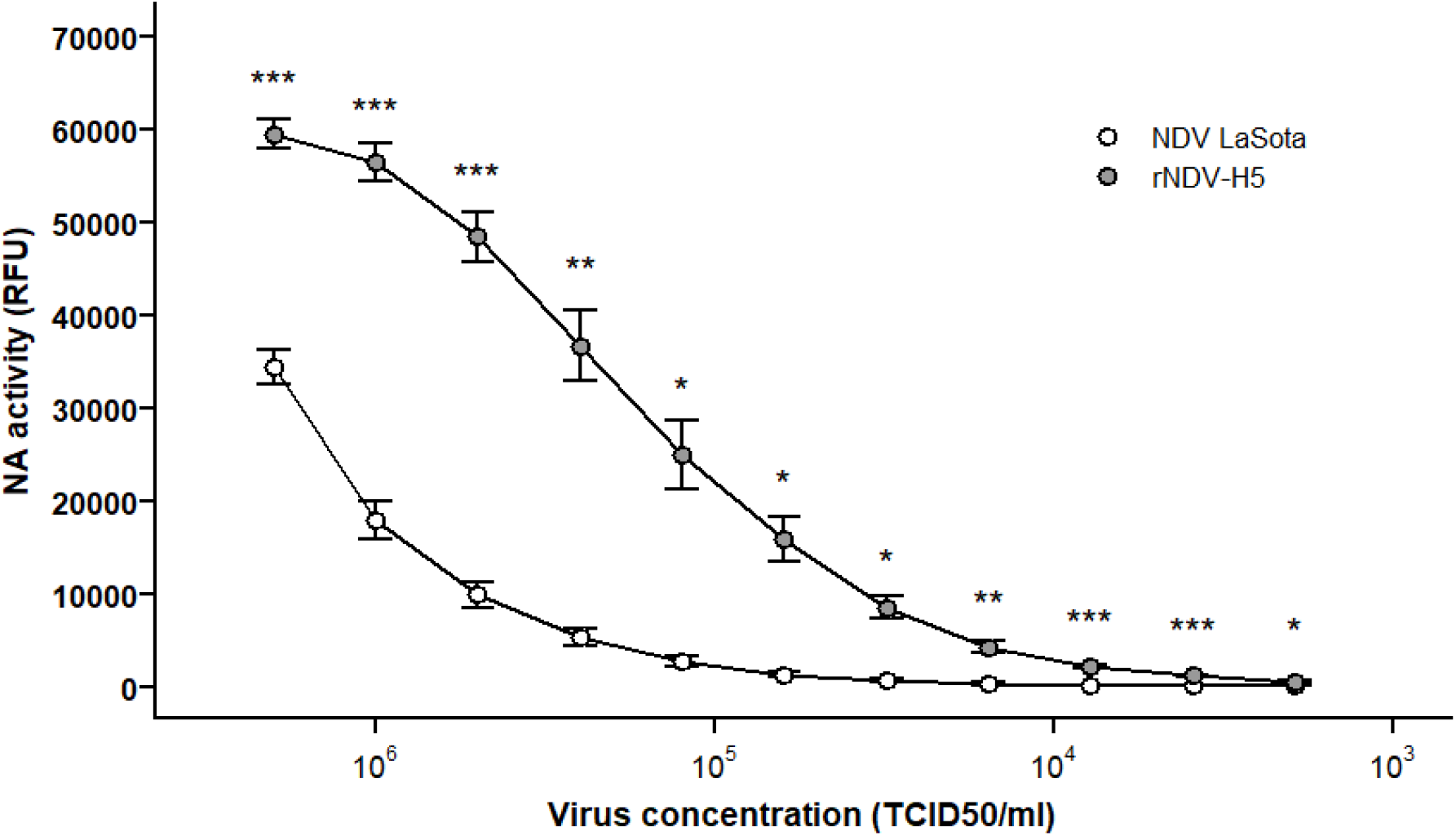
Hemagglutinating and neuraminidase activities of NDV LaSota and rNDV-H5. (a) Hemagglutination of chicken erythrocytes by two-fold serially diluted NDV LaSota (10^7^ TCID50/ml) and rNDV-H5 (10^7^ TCID50/ml). NC, negative control. (b) Neuraminidase activity of two-fold serially diluted NDV LaSota and rNDV-H5 viruses starting at a titer of 10^7^ TCID50/ml.* p < 0.05, ** p < 0.01, ***,p < 0.001.

The comparison of the NA activity showed that enzymatic activity of rNDV-H5 was significantly increased relative to that of the parental NDV LaSota at all virus titers tested (**Figure 2b**), which is in accordance with the higher HN content of rNDV-H5.

### Innate sensing of rNDV-H5 is mediated by TLR3, MDA5, and LGP2

CEFs were infected in triplicates with either NDV LaSota or rNDV-H5 at an MOI of 1. Representative images are shown in **Figure 3a**. CEFs infected with NDV LaSota and rNDV-H5 displayed a similar pattern of immunofluorescence for the F glycoprotein. However, infection kinetics differed slightly between both viruses, as NDV LaSota infection was more notable at 10 hpi than rNDV-H5.

**Figure 3.**
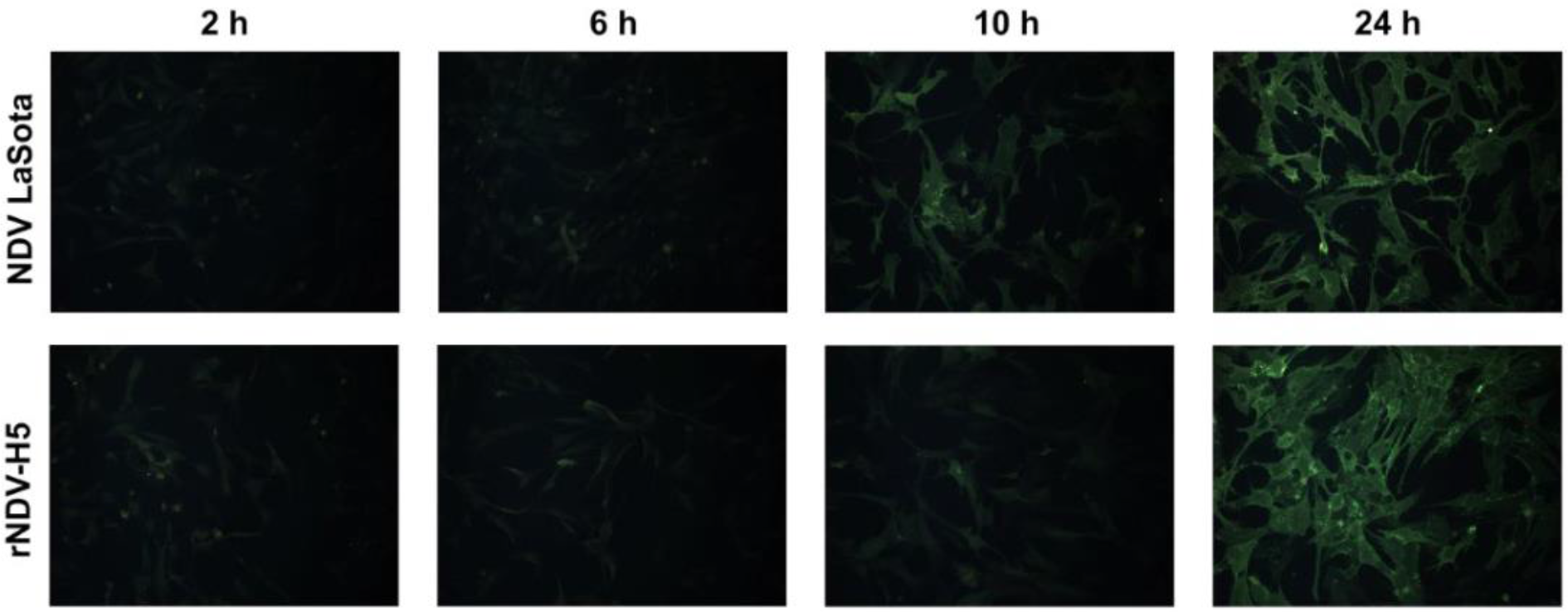
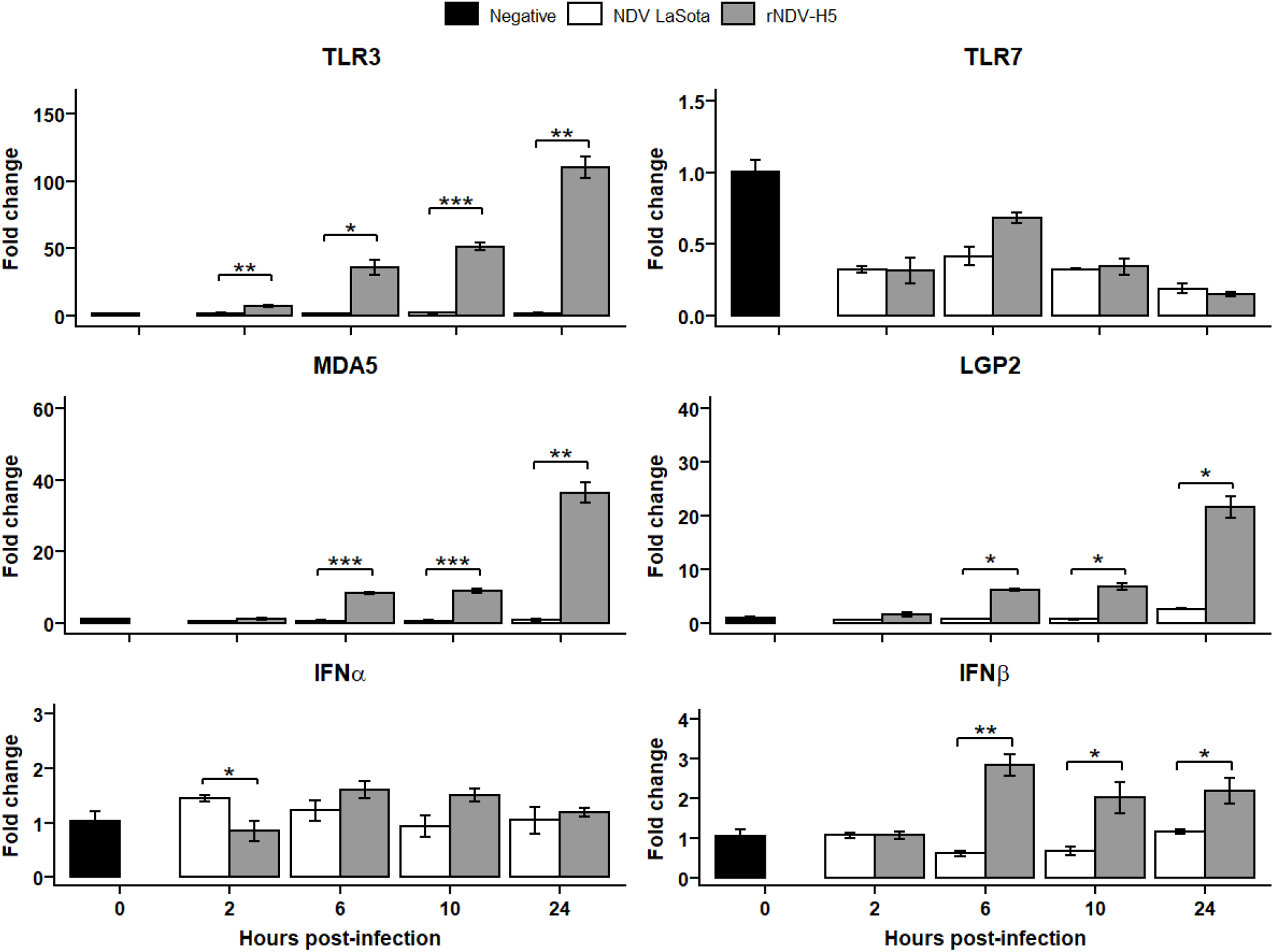
PRRs and type-I IFNs expression in NDV LaSota- and rNDV-H5-infected CEFs. (a) CEFs were infected with NDV LaSota and rNDV-H5 (MOI = 1) and observed by fluorescence microscopy using an anti-F antibody (IC3, IgG1) at 2, 6, 10, and 24 hpi. (b) Relative expression of TLR3, TLR7, MDA5, LGP2, IFNα, and IFNβ was determined in CEFs infected with NDV LaSota and rNDV-H5 (MOI = 1) at 2, 6, 10, and 24 hpi. The data were normalized to HMBS and RPL0 expression, calculated according to the 2^−ΔΔCT^ method, and presented ± standard error of the mean. * p < 0.05, ** p < 0.01, ***,p < 0.001.

Early immune responses induced following the infection of CEFs (**Figure 3b**) and TOCs (**Figure 4**) with rNDV-H5 and NDV LaSota were investigated through the evaluation of changes in the expression of genes associated with innate immune responses. The innate sensing of rNDV-H5 and NDV LaSota was first evaluated through the investigation of changes in the expression of PRRs TLR3, TLR7, MDA5, and LGP2.

**Figure 4.**
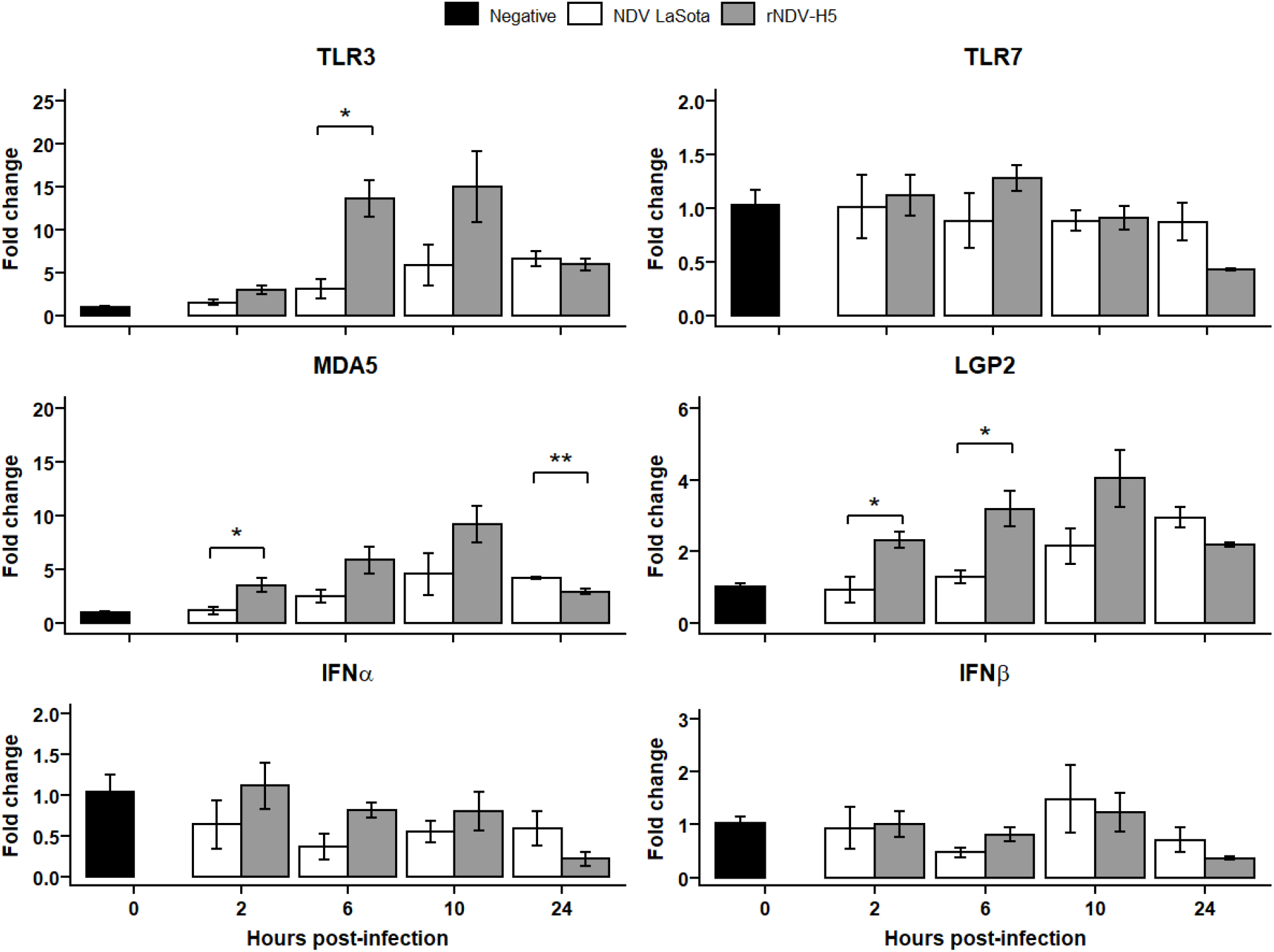
PRRs and type-I IFNs expression in NDV LaSota- and rNDV-H5-infected TOCs. Relative expression of TLR3, TLR7, MDA5, LGP2, IFNα, and IFNβ was determined in TOCs infected with NDV LaSota and rNDV-H5 (MOI = 1) at 2, 6, 10, and 24 hpi. The data were normalized to HMBS and RPL0 expression, calculated according to the 2^−ΔΔCT^ method, and presented ± standard error of the mean. * p < 0.05, ** p < 0.01, ***,p < 0.001.

A significantly increased expression of TLR3 at 2, 6, 10, and 24 hpi was observed in rNDV-H5 infections of CEFs, as compared with NDV LaSota. Expression of TLR7 was not significantly altered, regardless of the condition of infection. An increase in the expression of MDA5 and LGP2 was detected in CEFs infected with rNDV-H5 at 6, 10, and 24 hpi, as compared with NDV LaSota-infected cells.

The expression of TLR3 was significantly increased in the TOCs infected with rNDV-H5 compared to those infected with NDV LaSota at 6 hpi. No differences were observed in TLR7 gene expression between rNDV-H5- and NDV LaSota-infected TOCs. At 2 hpi, rNDV-H5-infected TOCs presented higher MDA5 gene expression than did those infected with NDV LaSota, and this trend was maintained at 6 and 10 hpi but was not significant. However, MDA5 expression was significantly higher in TOCs infected with NDV LaSota at 24 hpi compared to rNDV-H5 infection. The expression of LGP2 by rNDV-H5-infected TOCs was moderately upregulated at 2 and 6 hpi.

The expression of antiviral type I IFNs, IFNα and IFNβ, was also evaluated to determine whether it correlated changes in PRRs expression. A relatively modest decrease in IFNα expression was observed in cells infected with rNDV-H5 compared to NDV LaSota CEFs infection. In contrast, expression levels of IFNβ were increased in CEFs infected with rNDV-H5 at 6, 10, and 24 hpi, compared with NDV LaSota infection. These changes mirrored the decreased expression levels of MDA5 and LGP2 at the same time point. None of the TOCs infections significantly affected the expression of antiviral IFNα and IFNβ, regardless of the tested time point.

## DISCUSSION

The structure of rNDV-H5 and the expression of H5 on the vaccine surface have previously been observed by immunogold electron microscopy [18]. The present study confirmed these results and found significant structural differences between rNDV-H5 and the parental LaSota NDV. Because H5 gene is inserted upstream of the 3’ end of the genome of the rNDV-H5 construct, between phosphoprotein and matrix genes, the level of expression of downstream F and HN could be impacted and reduced [43,44]. However, the investigation of both surface glycoproteins distribution demonstrated a higher expression of HN and a lower expression of F by rNDV-H5 when compared to parental NDV LaSota, although HN is positioned more distally than F in the NDV genome. Earlier studies have shown that HN interacts with F and promotes its fusion activity [45,46]. During budding and virion assembly processes, HN and F are anchored into the viral envelope by the interaction of their cytoplasmic tail with the M protein [47]. The HA at the surface of rNDV-H5 is expressed as a chimeric protein whose H5 transmembrane domain and cytoplasmic tail have been replaced by those of the NDV F glycoprotein. Nayak et al. demonstrated that chimeric H5 was incorporated more efficiently into the NDV when compared to unmodified H5, but reduced replication of the recombinant vector without impacting its pathogenicity, suggesting that H5 incorporation may have affected the NDV assembly process [11]. The slight delay in the infection kinetics of CEFs with rNDV-H5 and the lower level of expression of F protein observed in the present study suggest that the incorporation of the chimeric HA might be at the expense of F incorporation. The structural impact of H5 incorporation could be confirmed by comparing the expression levels of F protein on the surface of recombinant NDV expressing chimeric or native HA.

Balanced hemagglutinin and neuraminidase enzyme activities are known to be critical for the outcome of an infection [45,48,49]. Following viral budding, the newly synthesized virus particles bound to the sialic acids of the host cells are released by the sialidase activity of NA (reviewed in [50]). A weak NA activity was found to be counterbalanced by a decreased HA affinity, maintaining the ability of the virus to replicate efficiently [51,52]. The results of the present study suggest that the presence of H5 on the surface confers a slightly higher hemagglutinating activity to rNDV-H5 compared to the parental LaSota NDV. In addition to the increased neuraminidase activity carried by rNDV-H5, these results raised the hypothesis that the presence of H5 presumably disrupted the HA:NA balance and may have been compensated by an enhanced neuraminidase activity as a result of a higher expression of HN protein. The comparison of the HN sequences of rNDV-H5 and LaSota NDV should be performed to evaluate the hypothesis that the increase in neuraminidase activity is indeed correlated to the increased number of HN proteins on the surface of the recombinant virus and not due to acquired compensatory mutations, as previously demonstrated for avian influenza in the case of an imbalance of hemagglutinating and NA activities [53].

Vaccination with rNDV-H5 has been previously demonstrated to offer enhanced protection against AI-infection to chickens carrying NDV-specific maternal antibodies, suggesting that viral entry into the host cell may have been partially H5-dependent [18]. In addition to these findings, the structural differences observed between rNDV-H5 and native NDV LaSota raised the question of their impact on the immune sensing of the recombinant virus by the host immune system. Elucidating the activation of PRRs by rNDV-H5 is fundamental for improving our understanding of protection outcomes previously demonstrated with these vaccines. The results of the present study demonstrated that the innate immune sensing of rNDV-H5 is mediated by TLR3, MDA5, and LGP2 receptors during early infection of CEFs and TOCs, resulting in the induction of IFNβ expression in CEFs. The activation and the antiviral effect of TLR3 during NDV LaSota infection have been previously observed in chicken embryo fibroblasts cell line overexpressing TLR3 [54]. The infection with rNDV-H5 induced a dramatic increase in TLR3 expression in comparison to NDV LaSota, suggesting that the expression of exogenous H5 by the vector modified the way it is recognized by the innate immunity sensors. This raised the hypothesis that reduced F expression and the presence of H5 might impact the entry pathway by partly promoting H5-dependent endocytosis during the early phase of infection, instead of the direct fusion entry predominantly used by NDV. Nevertheless, rNDV-H5 retains the ability to induce cytoplasmic PRRs MDA5 and LGP2, demonstrating the presence of genetic material in the cytoplasm. The expression of TLR7, which also senses viral RNA within endosomal compartments, has been previously detected in B cell-like DT40 and macrophage-like HD11 cell lines but not in CEFs [55]. Moreover, the expression of TLR7 previously examined in NDV LaSota-infected HD11 was not found to be altered [56]. The latter finding is in accordance with the lack of detection of TLR7 activation observed in the present study in CEFs and TOCs after NDV LaSota or rNDV-H5 infection. Finally, Paramyxoviruses use a mechanism involving the V protein to evade the host’s innate immune responses. The V protein is produced by RNA editing of the phosphoprotein (P) gene [57]. The V protein antagonizes the induction of type-I IFNs by binding both MDA5 and LGP2 [58–60], which was demonstrated to be correlated with NDV virulence [61]. V protein expression could be investigated to evaluate whether the insertion of H5 downstream of P may have influenced the expression of V by rNDV-H5 and contribute to the increased expression of IFNβ during cell infection by rNDV-H5.

Overall, this study demonstrated that the expression of a recombinant H5 by a recombinant NDV can not only impact its biological and structural characteristics but also induce changes in the recognition of the vector by innate immunity. These results confirm the need to systematically investigate the impact of the expression of a foreign gene by commonly used vaccine vectors, such as NDV, in order to improve the understanding of the protection against strains associated with the foreign gene and the vector itself.

## ABBREVIATIONS

CEFs: chicken embryo fibroblasts
F: fusion protein
HA: hemagglutinin
HN: hemagglutinin-neuraminidase protein
HPAI: highly pathogenic avian influenza
hpi: hours post-infection
IFNs: interferons
LGP2: laboratory of genetics and physiology 2
mAbs: monoclonal antibodies
MDA5: melanoma differentiation-associated gene 5
MOI: multiplicity of infection
NA: neuraminidase
NDV: Newcastle disease virus
PRRs: pattern recognition receptors
rNDV: recombinant NDV
RT: room temperature
SPF: specific pathogen free
TLRs: toll-like receptors
TOCs: tracheal organ cultures

## ACKNOWLEDGMENTS

This work was financially supported by the BELVIR-consortium (FNRS, Belgium) and Sciensano. The authors thank Alexandre Ausloos, Marc Boschmans, Eva Ngabirano, and Catherine Rasseneur, members of the Sciensano’s ‘avian virology and immunology’ team, and Marina Ledecq of the Sciensano’s ‘trace elements and nanomaterials’ service for their help and support in the realization of the experiments. The authors wish to express their thanks to Lohmann Animal Health GmbH (Germany) for providing the rNDV-H5 and the NDV LaSota vaccines.

## REFERENCES

1. Astill, J.; Dara, R.A.; Fraser, E.D.G.; Sharif, S. Detecting and Predicting Emerging Disease in Poultry With the Implementation of New Technologies and Big Data: A Focus on Avian Influenza Virus. Front. Vet. Sci. 2018, 5, doi:10.3389/fvets.2018.00263.

2. Bello, M.B.; Yusoff, K.; Ideris, A.; Hair-Bejo, M.; Peeters, B.P.H.; Omar, A.R. Diagnostic and Vaccination Approaches for Newcastle Disease Virus in Poultry: The Current and Emerging Perspectives. BioMed Res. Int. 2018, 2018, 1–18, doi:10.1155/2018/7278459.

3. Suarez, D.L.; Pantin-Jackwood, M.J. Recombinant Viral-Vectored Vaccines for the Control of Avian Influenza in Poultry. Vet. Microbiol. 2017, 206, 144–151, doi:10.1016/j.vetmic.2016.11.025.

4. Dimitrov, K.M.; Afonso, C.L.; Yu, Q.; Miller, P.J. Newcastle Disease Vaccines—A Solved Problem or a Continuous Challenge? Vet. Microbiol. 2017, 206, 126–136, doi:10.1016/j.vetmic.2016.12.019.

5. Spackman, E.; Pantin-Jackwood, M.J. Practical Aspects of Vaccination of Poultry against Avian Influenza Virus. Vet. J. 2014, 202, 408–415, doi:10.1016/j.tvjl.2014.09.017.

6. Romanutti, C.; Keller, L.; Zanetti, F.A. Current Status of Virus-Vectored Vaccines against Pathogens That Affect Poultry. Vaccine 2020, 38, 6990–7001, doi:10.1016/j.vaccine.2020.09.013.

7. Kim, S.-H.; Samal, S.K. Newcastle Disease Virus as a Vaccine Vector for Development of Human and Veterinary Vaccines. Viruses 2016, 8, doi:10.3390/v8070183.

8. Ge, J.; Deng, G.; Wen, Z.; Tian, G.; Wang, Y.; Shi, J.; Wang, X.; Li, Y.; Hu, S.; Jiang, Y.; et al. Newcastle Disease Virus-Based Live Attenuated Vaccine Completely Protects Chickens and Mice from Lethal Challenge of Homologous and Heterologous H5N1 Avian Influenza Viruses. J. Virol. 2007, 81, 150–158, doi:10.1128/JVI.01514-06.

9. Kim, S.-H.; Samal, S.K. Innovation in Newcastle Disease Virus Vectored Avian Influenza Vaccines. Viruses 2019, 11, doi:10.3390/v11030300.

10. Lardinois, A.; Steensels, M.; Lambrecht, B.; Desloges, N.; Rahaus, M.; Rebeski, D.; van den Berg, T. Potency of a Recombinant NDV-H5 Vaccine Against Various HPAI H5N1 Virus Challenges in SPF Chickens. Avian Dis. 2012, 56, 928–936, doi:10.1637/10173-041012-ResNote.1.

11. Nayak, B.; Rout, S.N.; Kumar, S.; Khalil, M.S.; Fouda, M.M.; Ahmed, L.E.; Earhart, K.C.; Perez, D.R.; Collins, P.L.; Samal, S.K. Immunization of Chickens with Newcastle Disease Virus Expressing H5 Hemagglutinin Protects against Highly Pathogenic H5N1 Avian Influenza Viruses. PLOS ONE 2009, 4, e6509, doi:10.1371/journal.pone.0006509.

12. Veits, J.; Wiesner, D.; Fuchs, W.; Hoffmann, B.; Granzow, H.; Starick, E.; Mundt, E.; Schirrmeier, H.; Mebatsion, T.; Mettenleiter, T.C.; et al. Newcastle Disease Virus Expressing H5 Hemagglutinin Gene Protects Chickens against Newcastle Disease and Avian Influenza. Proc. Natl. Acad. Sci. 2006, 103, 8197–8202, doi:10.1073/pnas.0602461103.

13. Connaris, H.; Takimoto, T.; Russell, R.; Crennell, S.; Moustafa, I.; Portner, A.; Taylor, G. Probing the Sialic Acid Binding Site of the Hemagglutinin-Neuraminidase of Newcastle Disease Virus: Identification of Key Amino Acids Involved in Cell Binding, Catalysis, and Fusion. J. Virol. 2002, 76, 1816–1824, doi:10.1128/JVI.76.4.1816-1824.2002.

14. Aguilar, H.C.; Henderson, B.A.; Zamora, J.L.; Johnston, G.P. Paramyxovirus Glycoproteins and the Membrane Fusion Process. Curr. Clin. Microbiol. Rep. 2016, 3, 142–154, doi:10.1007/s40588-016-0040-8.

15. Cantin, C.; Holguera, J.; Ferreira, L.; Villar, E.; Munoz-Barroso, I. Newcastle Disease Virus May Enter Cells by Caveolae-Mediated Endocytosis. J. Gen. Virol. 2007, 88, 559–569, doi:10.1099/vir.0.82150-0.

16. Wen, Z.; Zhao, B.; Song, K.; Hu, X.; Chen, W.; Kong, D.; Ge, J.; Bu, Z. Recombinant Lentogenic Newcastle Disease Virus Expressing Ebola Virus GP Infects Cells Independently of Exogenous Trypsin and Uses Macropinocytosis as the Major Pathway for Cell Entry. Virol. J. 2013, 10, 331, doi:10.1186/1743-422X-10-331.

17. Edinger, T.O.; Pohl, M.O.; Stertz, S. Entry of Influenza A Virus: Host Factors and Antiviral Targets. J. Gen. Virol. 2014, 95, 263–277, doi:10.1099/vir.0.059477-0.

18. Lardinois, A.; Vandersleyen, O.; Steensels, M.; Desloges, N.; Mast, J.; van den Berg, T.; Lambrecht, B. Stronger Interference of Avian Influenza Virus–Specific Than Newcastle Disease Virus–Specific Maternally Derived Antibodies with a Recombinant NDV-H5 Vaccine. Avian Dis. 2016, 60, 191–201, doi:10.1637/11133-050815-Reg.

19. Locati, M.; Mantovani, A.; Sica, A. Macrophage Activation and Polarization as an Adaptive Component of Innate Immunity. Adv. Immunol. 2013, 120, 163–184, doi:10.1016/B978-0-12-417028-5.00006-5.

20. O’Neill, L.A.J.; Bowie, A.G. Sensing and Signaling in Antiviral Innate Immunity. Curr. Biol. 2010, 20, R328–R333, doi:10.1016/j.cub.2010.01.044.

21. Chen, S.; Cheng, A.; Wang, M. Innate Sensing of Viruses by Pattern Recognition Receptors in Birds. Vet. Res. 2013, 44, 82, doi:10.1186/1297-9716-44-82.

22. Denney, L.; Ho, L.-P. The Role of Respiratory Epithelium in Host Defence against Influenza Virus Infection. Biomed. J. 2018, 41, 218–233, doi:10.1016/j.bj.2018.08.004.

23. Santhakumar, D.; Rubbenstroth, D.; Martinez-Sobrido, L.; Munir, M. Avian Interferons and Their Antiviral Effectors. Front. Immunol. 2017, 8, doi:10.3389/fimmu.2017.00049.

24. Park, M.-S.; Steel, J.; García-Sastre, A.; Swayne, D.; Palese, P. Engineered Viral Vaccine Constructs with Dual Specificity: Avian Influenza and Newcastle Disease. Proc. Natl. Acad. Sci. 2006, 103, 8203–8208, doi:10.1073/pnas.0602566103.

25. Zhao, W.; Zhang, Z.; Zsak, L.; Yu, Q. P and M Gene Junction Is the Optimal Insertion Site in Newcastle Disease Virus Vaccine Vector for Foreign Gene Expression. J. Gen. Virol. 2015, 96, 40–45, doi:10.1099/vir.0.068437-0.

26. Steensels, M.; Van Borm, S.; Boschmans, M.; van den Berg, T. Lethality and Molecular Characterization of an HPAI H5N1 Virus Isolated from Eagles Smuggled from Thailand into Europe. Avian Dis. 2007, 51, 401–407, doi:10.1637/7554-033106R.1.

27. Nagy, A.; Lee, J.; Mena, I.; Henningson, J.; Li, Y.; Ma, J.; Duff, M.; Li, Y.; Lang, Y.; Yang, J.; et al. Recombinant Newcastle Disease Virus Expressing H9 HA Protects Chickens against Heterologous Avian Influenza H9N2 Virus Challenge. Vaccine 2016, 34, 2537–2545, doi:10.1016/j.vaccine.2016.04.022.

28. Shahsavandi, S.; Ebrahimi, M.M.; Mohammadi, A.; Zarrin Lebas, N. Impact of Chicken-Origin Cells on Adaptation of a Low Pathogenic Influenza Virus. Cytotechnology 2013, 65, 419–424, doi:10.1007/s10616-012-9495-5.

29. Ren, X.; Xue, C.; Kong, Q.; Zhang, C.; Bi, Y.; Cao, Y. Proteomic Analysis of Purified Newcastle Disease Virus Particles. Proteome Sci. 2012, 10, 32, doi:10.1186/1477-5956-10-32.

30. Ferreira, H.L.; Lambrecht, B.; van Borm, S.; Torrieri-Dramard, L.; Klatzmann, D.; Bellier, B.; van den Berg, T. Identification of a Dominant Epitope in the Hemagglutinin of an Asian Highly Pathogenic Avian Influenza H5N1 Clade 1 Virus by Selection of Escape Mutants. Avian Dis. 2010, 54, 565–571, doi:10.1637/8750-033009-ResNote.1.

31. Meulemans, G.; Gonze, M.; Carlier, M.C.; Petit, P.; Burny, A.; Le Long, null Evaluation of the Use of Monoclonal Antibodies to Hemagglutinin and Fusion Glycoproteins of Newcastle Disease Virus for Virus Identification and Strain Differentiation Purposes. Arch. Virol. 1987, 92, 55–62, doi:10.1007/BF01310062.

32. Kortekaas, J.; Dekker, A.; de Boer, S.M.; Weerdmeester, K.; Vloet, R.P.M.; Wit, A.A.C. de; Peeters, B.P.H.; Moormann, R.J.M. Intramuscular Inoculation of Calves with an Experimental Newcastle Disease Virus-Based Vector Vaccine Elicits Neutralizing Antibodies against Rift Valley Fever Virus. Vaccine 2010, 28, 2271–2276, doi:10.1016/j.vaccine.2010.01.001.

33. Spackman, E. Animal Influenza Virus; Second edition.; Humana Press: New York, 2014; ISBN 978-1-4939-0757-1.

34. Ingrao, F.; Rauw, F.; van den Berg, T.; Lambrecht, B. Characterization of Two Recombinant HVT-IBD Vaccines by VP2 Insert Detection and Cell-Mediated Immunity after Vaccination of Specific Pathogen-Free Chickens. Avian Pathol. 2017, 46, 289–299, doi:10.1080/03079457.2016.1265083.

35. Wang, J.; Tang, C.; Wang, Q.; Li, R.; Chen, Z.; Han, X.; Wang, J.; Xu, X. Apoptosis Induction and Release of Inflammatory Cytokines in the Oviduct of Egg-Laying Hens Experimentally Infected with H9N2 Avian Influenza Virus. Vet. Microbiol. 2015, 177, 302–314, doi:10.1016/j.vetmic.2015.04.005.

36. He, Y.; Xie, Z.; Dai, J.; Cao, Y.; Hou, J.; Zheng, Y.; Wei, T.; Mo, M.; Wei, P. Responses of the Toll-like Receptor and Melanoma Differentiation-Associated Protein 5 Signaling Pathways to Avian Infectious Bronchitis Virus Infection in Chicks. Virol. Sin. 2016, 31, 57–68, doi:10.1007/s12250-015-3696-y.

37. Peters, M.A.; Browning, G.F.; Washington, E.A.; Crabb, B.S.; Kaiser, P. Embryonic Age Influences the Capacity for Cytokine Induction in Chicken Thymocytes. Immunology 2003, 110, 358–367, doi:https://doi.org/10.1046/j.1365-2567.2003.01744.x.

38. Nang, N.T.; Lee, J.S.; Song, B.M.; Kang, Y.M.; Kim, H.S.; Seo, S.H. Induction of Inflammatory Cytokines and Toll-like Receptors in Chickens Infected with Avian H9N2 Influenza Virus. Vet. Res. 2011, 42, 64, doi:10.1186/1297-9716-42-64.

39. Ingrao, F.; Rauw, F.; Steensels, M.; van den Berg, T.; Lambrecht, B. Early Immune Responses and Profiling of Cell-Mediated Immunity-Associated Gene Expression in Response to RHVT-IBD Vaccination. Vaccine 2018, 36, 615–623, doi:10.1016/j.vaccine.2017.12.059.

40. Staines, K.; Batra, A.; Mwangi, W.; Maier, H.J.; Van Borm, S.; Young, J.R.; Fife, M.; Butter, C. A Versatile Panel of Reference Gene Assays for the Measurement of Chicken MRNA by Quantitative PCR. PLOS ONE 2016, 11, e0160173, doi:10.1371/journal.pone.0160173.

41. Livak, K.J.; Schmittgen, T.D. Analysis of Relative Gene Expression Data Using Real-Time Quantitative PCR and the 2−ΔΔCT Method. Methods 2001, 25, 402–408, doi:10.1006/meth.2001.1262.

42. Wickham, H. Ggplot2: Elegant Graphics for Data Analysis; Use R!; 2nd ed.; Springer International Publishing, 2016; ISBN 978-3-319-24275-0.

43. Skiadopoulos, M.H.; Surman, S.R.; Riggs, J.M.; Örvell, C.; Collins, P.L.; Murphy, B.R. Evaluation of the Replication and Immunogenicity of Recombinant Human Parainfluenza Virus Type 3 Vectors Expressing up to Three Foreign Glycoproteins. Virology 2002, 297, 136–152, doi:10.1006/viro.2002.1415.

44. Willemsen, A.; Zwart, M.P. On the Stability of Sequences Inserted into Viral Genomes. Virus Evol. 2019, 5, doi:10.1093/ve/vez045.

45. Jin, J.; Zhao, J.; Ren, Y.; Zhong, Q.; Zhang, G. Contribution of HN Protein Length Diversity to Newcastle Disease Virus Virulence, Replication and Biological Activities. Sci. Rep. 2016, 6, 36890, doi:10.1038/srep36890.

46. Melanson, V.R.; Iorio, R.M. Amino Acid Substitutions in the F-Specific Domain in the Stalk of the Newcastle Disease Virus HN Protein Modulate Fusion and Interfere with Its Interaction with the F Protein. J. Virol. 2004, 78, 13053–13061, doi:10.1128/JVI.78.23.13053-13061.2004.

47. Liu, Y.C.; Grusovin, J.; Adams, T.E. Electrostatic Interactions between Hendra Virus Matrix Proteins Are Required for Efficient Virus-Like-Particle Assembly. J. Virol. 2018, 92, doi:10.1128/JVI.00143-18.

48. Tappert, M.M.; Porterfield, J.Z.; Mehta-D’Souza, P.; Gulati, S.; Air, G.M. Quantitative Comparison of Human Parainfluenza Virus Hemagglutinin-Neuraminidase Receptor Binding and Receptor Cleavage. J. Virol. 2013, 87, 8962–8970, doi:10.1128/JVI.00739-13.

49. Liu, T.; Song, Y.; Yang, Y.; Bu, Y.; Cheng, J.; Zhang, G.; Xue, J. Hemagglutinin– Neuraminidase and Fusion Genes Are Determinants of NDV Thermostability. Vet. Microbiol. 2019, 228, 53–60, doi:10.1016/j.vetmic.2018.11.013.

50. Bouvier, N.M.; Palese, P. The Biology of Influenza Viruses. Vaccine 2008, 26, D49–D53, doi:10.1016/j.vaccine.2008.07.039.

51. Richard, M.; Ferraris, O.; Erny, A.; Barthélémy, M.; Traversier, A.; Sabatier, M.; Hay, A.; Lin, Y.P.; Russell, R.J.; Lina, B. Combinatorial Effect of Two Framework Mutations (E119V and I222L) in the Neuraminidase Active Site of H3N2 Influenza Virus on Resistance to Oseltamivir. Antimicrob. Agents Chemother. 2011, 55, 2942–2952, doi:10.1128/AAC.01699-10.

52. Xu, R.; Zhu, X.; McBride, R.; Nycholat, C.M.; Yu, W.; Paulson, J.C.; Wilson, I.A. Functional Balance of the Hemagglutinin and Neuraminidase Activities Accompanies the Emergence of the 2009 H1N1 Influenza Pandemic. J. Virol. 2012, 86, 9221–9232, doi:10.1128/JVI.00697-12.

53. Du, W.; Wolfert, M.A.; Peeters, B.; Kuppeveld, F.J.M. van; Boons, G.-J.; Vries, E. de; Haan, C.A.M. de Mutation of the Second Sialic Acid-Binding Site of Influenza A Virus Neuraminidase Drives Compensatory Mutations in Hemagglutinin. PLOS Pathog. 2020, 16, e1008816, doi:10.1371/journal.ppat.1008816.

54. Cheng, J.; Sun, Y.; Zhang, X.; Zhang, F.; Zhang, S.; Yu, S.; Qiu, X.; Tan, L.; Song, C.; Gao, S.; et al. Toll-like Receptor 3 Inhibits Newcastle Disease Virus Replication through Activation of pro-Inflammatory Cytokines and the Type-1 Interferon Pathway. Arch. Virol. 2014, 159, 2937–2948, doi:10.1007/s00705-014-2148-6.

55. Philbin, V.J.; Iqbal, M.; Boyd, Y.; Goodchild, M.J.; Beal, R.K.; Bumstead, N.; Young, J.; Smith, A.L. Identification and Characterization of a Functional, Alternatively Spliced Toll-like Receptor 7 (TLR7) and Genomic Disruption of TLR8 in Chickens. Immunology 2005, 114, 507–521, doi:https://doi.org/10.1111/j.1365-2567.2005.02125.x.

56. Zhang, P.; Ding, Z.; Liu, X.; Chen, Y.; Li, J.; Tao, Z.; Fei, Y.; Xue, C.; Qian, J.; Wang, X.; et al. Enhanced Replication of Virulent Newcastle Disease Virus in Chicken Macrophages Is Due to Polarized Activation of Cells by Inhibition of TLR7. Front. Immunol. 2018, 9, doi:10.3389/fimmu.2018.00366.

57. Steward, M.; Vipond, I.B.; Millar, N.S.; Emmerson, P.T. RNA Editing in Newcastle Disease Virus. J. Gen. Virol. 1993, 74, 2539–2547, doi:10.1099/0022-1317-74-12-2539.

58. Childs, K.; Stock, N.; Ross, C.; Andrejeva, J.; Hilton, L.; Skinner, M.; Randall, R.; Goodbourn, S. Mda-5, but Not RIG-I, Is a Common Target for Paramyxovirus V Proteins. Virology 2007, 359, 190–200, doi:10.1016/j.virol.2006.09.023.

59. Childs, K.; Randall, R.; Goodbourn, S. Paramyxovirus V Proteins Interact with the RNA Helicase LGP2 To Inhibit RIG-I-Dependent Interferon Induction. J. Virol. 2012, 86, 3411–3421, doi:10.1128/JVI.06405-11.

60. Childs, K.S.; Randall, R.E.; Goodbourn, S. LGP2 Plays a Critical Role in Sensitizing Mda-5 to Activation by Double-Stranded RNA. PLoS ONE 2013, 8, doi:10.1371/journal.pone.0064202.

61. Wang, X.; Dang, R.; Yang, Z. The Interferon Antagonistic Activities of the V Proteins of NDV Correlated with Their Virulence. Virus Genes 2019, 55, 233–237, doi:10.1007/s11262-019-01637-3.

